# Adaptive evolution of transcriptome and transcriptomic plasticity in stable and fluctuating environments

**DOI:** 10.64898/2026.02.16.706080

**Authors:** Arnaud Le Rouzic, Apolline Petit, Arthur Jallet, Justine Vergne, Ekta Kochar, Anne Genissel

## Abstract

Adaptation to new environments often involves multiple genes and can induce substantial changes in biological systems, but the overall consequences of the genetic response to specific environmental pressures remain poorly understood. Here, we investigate the large-scale effects of adaptation to new temperature regimes on the whole transcriptome from experimental evolution carried on the fungus *Zymoseptoria tritici*. Two distinct strains from the wild were grown in constant cool (17°C), constant warm (23°C), and fluctuating temperature environments, for about 250 clonal generations. The expression of > 10,000 genes was estimated by RNA-seq before and after evolution at several temperatures. We observed massive convergent change in both gene expression and gene expression plasticity. Fluctuating environments did not favor plastic gene expression in general, although fluctuating populations did evolve towards the optimal reaction norms. Some of our observations were at odds with common expectations; adaptive mutations were highly pleiotropic, and adaptation to stable temperature conditions did not match gene expression plasticity. We thus showed that the evolution of complex biological systems follow some general patterns which challenge gene-centered or quantitative genetics predictions.

## Introduction

Genetic adaptation involves mutations that occur independently from the environment; mutations are sorted out by natural selection and genetic variants improving individual fitness invade the populations. Plastic adaptation, in contrast, relies on physiological, developmental, or behavioral responses to the environment; it occurs during the lifetime of individuals, and as such, does not participate to the evolution of species. It was early noticed that phenotypic plasticity itself was likely to be the consequence of genetic adaptation favoring plastic genotypes (Schmalhausen 1949; Bradshaw 1965), but the extent to which plasticity is at the core of the evolutionary process is still controversial (see Laland et al. 2014; Lewens 2019; Sommer 2020, for a summary of the arguments).

Theoretical quantitative genetics provide expectations about the circumstances in which plasticity should evolve as an adaptation (Via and Lande 1985; Lande 2009; Chevin et al. 2010). Three conditions have to be fulfilled: (i) plasticity can evolve (i.e., there must be genotype-by-environment interactions); (ii) the environmental conditions favor plastic genotypes (the optimal phenotype must be different in different environments, and the environment should change on a regular basis, so that populations can experience the fitness consequences of plasticity); (iii) organisms are able to perceive or predict the environment, either directly or indirectly. When all these conditions are not met simultaneously, plasticity could either stay constant, evolve as a consequence of genetic drift or correlated selection (non-adaptive plasticity), or selection could favor environmental robustness, i.e. the absence of plasticity. However, models generally focus on one or very few traits that are directly targeted by selection. In spite of a growing awareness that plasticity should be approached at the system level and not trait by trait (e.g., Spitze and Sadler 1996; Pennacchi et al. 2021), classical theory says little about the consequences of massively multivariate plasticity. Whether the simultaneous response of thousands of genes to environmental stimuli could be interpreted as the result of a coherent optimization process remains questionable, and cannot be answered based on theoretical models alone.

It is generally difficult to measure and understand selection and adaptation at the gene expression level (Fay and Wittkopp 2008; Price et al. 2022). A promising approach to accumulate empirical data about how gene expression evolves consists in sequencing mRNA during adaptation, typically in an experimental evolution setting (“evolve and resequence”) (Garland and Rose 2009; Kawecki et al. 2012; Long et al. 2015; Van den Bergh et al. 2018; Koch et al. 2025). Most of the time, experimental evolution tracks evolution in new, but stable, environmental conditions (Lenski et al. 1991), but adaptation to fluctuating environments has also motivated a series of historical experiments in bacteria (Leroi et al. 1994; Hughes et al. 2007; Ketola and Saarinen 2015). Similar experiments in unicellular eukaryotes (with their presumably more complex regulation system) were run more recently, with for instance alternated salt or oxidative stress in the yeast *Saccharomyces cerevisiae* (Dhar et al. 2013), or salinity fluctuations in the algae *Dunaliella salina* (Leung et al. 2020). Nevertheless, how gene regulatory systems adapt to fluctuating environments remains unclear: does gene expression become more sensitive to environmental change in unpredictable environments? Is gene expression plasticity lost in stable conditions? Does evolution tend to reinforce pre-existing plasticity? Is evolution of gene expression restricted to specific genes or modules, or is it a transcriptome-wide change?

Here, we propose to track the evolution of gene expression and gene expression plasticity during adaptation to both stable and fluctuating temperature conditions. Based on the experiment by Jallet et al. (2020), in which 17 populations of the wheat pathogen fungus *Zymoseptoria triciti* evolved for one year in lab conditions, we expanded and reanalyzed gene expression data at both 17°C and 23°C before and after evolution to assess the repeatability and convergence in gene expression and in gene expression plasticity in both stable and fluctuating lineages. Our focus here was not to identify genes and functions responding to selection, but rather to address the evolution of the transcriptome as a system. We considered convergent evolution across evolutionary replicates as a signature for adaptive evolution. More specifically, we aimed at (i) understanding the scale at which transcriptomes change as a response to different environmental pressures, (ii) measuring the consequences of adaptation to stable vs. fluctuating environments, and (iii) understanding how gene expression plasticity evolves as a response to these selection pressures.

## Materials and Methods

### Biological material and experimental evolution

This study is based on the experiment described in Jallet et al. (2020). While previous investigation focused at differentially-expressed genes associated with fluctuations from eight evolutionary lineages, the current work expanded this dataset with the transcriptome of 9 additional lineages (totalling 17), allowing the exploration of repeatability and evolutionary convergence in both stable and fluctuating lineages. All RNAseq data was reanalysed and mapped against a recent annotation.

Two clones of *Zymoseptoria triciti*, denoted as MGGP01 and MGGP44 were isolated in 2010 from a wheat field sample in Auzeville-Tolosane, France. Clones were acclimated to lab conditions (liquid medium: Potato Dextrose Broth – PDB – agitated at 140 rpm, constant temperature: 17°C) for a few weeks before being frozen in a PDB-glycerol mix at −80°C. In such lab conditions, *Z. tritici* grows mitotically, forming small (2 – 5 cells) structures distributed homogeneously in the liquid medium.

Experimental evolution took place for one year in 2015 and 2016. Both clonal initial populations were grown in agitated liquid PDB flasks to initiate 18 evolutionary lineages, submitted to 3 selection regimes: stable cool (17°C), stable warm (23°C), and fluctuating (17°C and 23°C alternating every 52-64 h) (Figure 1). Every week, 20mL of cell suspension (10^7^ to 10^9^ cells) were transferred into 500mL fresh medium, ensuring a weekly 26-fold amplification (about 4.7 mitotic divisions, totalling at least ∼225 generations over the 48 weeks of the experiment). Three evolutionary lineages were run for each selection regime and for each initial clone. By the end of the experiment, large volumes of cell suspensions were collected and frozen in 30% glycerol at −80°C.

**Figure 1.**
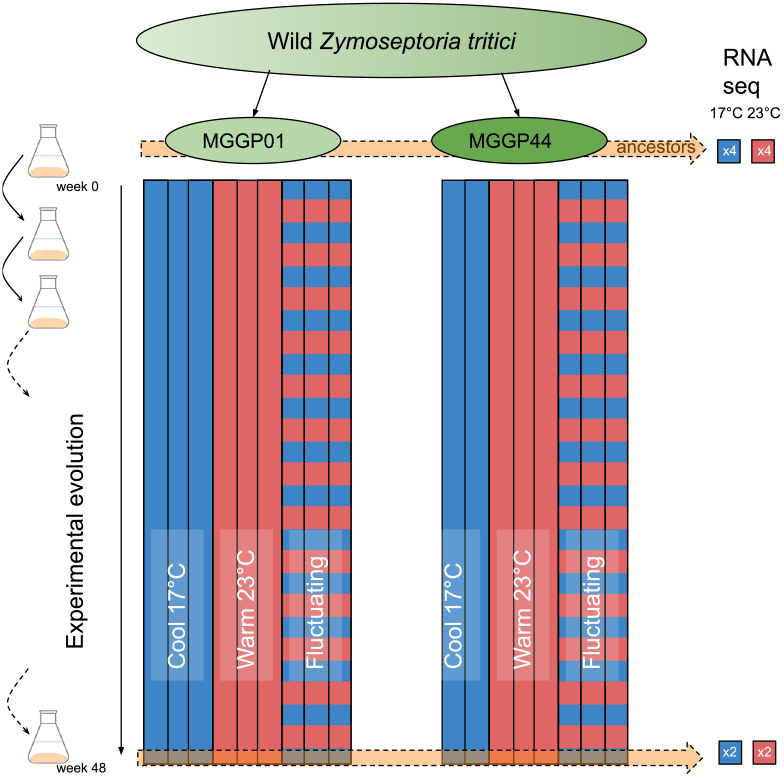
Experimental evolution setting. A total of 17 lineages were derived from two wild isolates of *Zymoseptoria tritici*, and submitted for 48 weeks to stable and fluctuating selection regimes (cool temperatures are represented in blue, warm temperatures in red). Note that one of the 17°C lineages was lost for genotype MGGP44 due to contamination, explaining the slight imbalance in the design. RNA was extracted and sequenced at both 17°C and 23°C for both ancestral genotypes, and after 48 weeks for evolved lineages. The experimental design involved 4 biological RNAseq replicates for ancestors at both temperatures, and 2 replicates for each of the evolved lines at both temperatures.

### RNA sequencing and bioinformatics

RNA was isolated after a 1-week growth stage at either 17°C or 23°C in the same conditions as for the experiment. Each strain (ancestors and evolved lineages) was grown and sequenced twice independently at each temperature, constituting pairs of biological replicates. The experimental procedure was expected to generate 2 genotypes × 3 selection regimes × 3 evolutionary replicates ×2 test temperatures × 2 replicates = 72 samples for the evolved populations, and 2 genotypes × 2 temperatures × 4 replicates = 16 samples for the parental populations. One of the 17°C lineage for MGGP44 appeared to be contaminated, and thus excluded from the analysis prior to sequencing, leaving 72 − 4 + 16 = 84 samples. For budget reasons, only about half (8/18) of the evolved lineages could be sequenced in 2016. The other 10 lineages were grown to produce and collect mycelium in the exact same conditions as for the first group in 2022, and were sequenced using the same technology. RNA was thus extracted and sequenced in 8 batches (4 at 17°C, 4 at 23°C; 4 in 2016 and 4 in 2022); pairs of biological replicates were always distributed in different batches, and both ancestral strains were present in each batch to serve as a reference when correcting for batch effects.

RNA was extracted as described extensively in Jallet et al. (2020). Cell suspension samples were snapfrozen, lyophilized for 3 days, and homogenized in a trisol - chloroform solution. Libraries were prepared with TruSeq Stranded mRNA Sample Prep kit from Illumina, and fragments between 250 and 400 bp were captured with poly-T oligonucleotides. Pair-end (2 × 75 bp) sequencing was performed with HiSeq4000 Illumina by Integragen Inc., Evry, France.

We acquired the read counts by following the pipeline described in Jallet et al. (2020), with the difference that we used a more recent *Z. tritici* annotation as reference (Lapalu et al. 2023). We used Trimmomatic (version 0.39) to remove adaptators and perform quality-based trimming. For each sample, mates of trimmed reads were mapped to the reference genome IPO-3232 by HISAT2 (version 2.2.1) using the parameters: `--min-intronlen 20 --max-intronlen 15,000 -k 3 --score-min L, 0, -0.4`. The quantification of mapped reads was done by Stringtie (version 2.2.1), with the options: `-p 16 -c 7 -m 200 -e {B -G`. Reads mapped to ribosomal units were removed from the analysis. In total, reads were mapped on 13,414 identified genes (12,729 for the MGGP01, and 12,871 for MGGP44).

### Data filtering and analysis

In order to perform a global quantitative analysis of gene expression, we focused on genes that were shared among both clones, and we excluded genes displaying 0 count in any sample (the vast majority of them being suspiciously inconsistent between pairs of biological replicates). Counts were normalized according to the Relative Log Expression method implemented in the R package DESeq2 (Love et al. 2014). Among the 84 samples, 2 were excluded because they have been either mislabelled or contaminated (labelled as MGGP44, but clearly belonging to MGGP01). Three other samples were also considered as outliers and excluded because they formed isolated clusters, far from their related lineages. The final dataset featured 10,641 genes and 79 samples, with 11 to 358,197 counts per gene (grand mean: 3257 counts per gene). Data analysis was performed with R version 4.1.2 (R Core Team 2021).

#### Quantitative analysis

Most of our analysis relies on a quantitative treatment of the gene expression data, considering log2 counts as quantitative traits. Batch effects were corrected by running the `removeBatchEffect` procedure from the limma package for R (Ritchie et al. 2015) independently for samples tested at 17°C and at 23°C.

We quantified the respective effect of experimental factors on the expression of all genes independently with the following linear model:

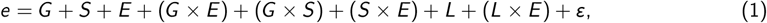

where *e* stands for the log2 of batch-effect corrected gene expression, *G* stands for the effect of genotype (2 levels: MGGP01 and MGGP44), *S* is the effect of the selection regime (4 levels: ancestor, stable at 17°C, stable at 23°C, and fluctuating), *E* quantifies the effect of plasticity (the effect of the environment in which gene expression was measured, 2 levels: 17°C and 23°C), *L* is the effect of the evolutionary lineage (17 evolved + 2 ancestors = 19 levels); *“* stands for the residuals (biological replicates). Interactions quantify the effect of experimental factors on plasticity: *G* × *E* is the difference in plasticity among genotypes, *G* × *S* is the differential response to selection depending on the genotype, *S* × *E* is the systematic effect of the selection regime on plasticity, and *L* × *E* the differences in plasticity among lineages. The lineage effect (evolutionary replicate) was treated as a random effect; genotype, temperature, and selection regime were fixed effects.

#### Convergent evolutio

Convergent evolution was quantified for each gene by a convergent evolution score that translates both the magnitude and the consistency of evolutionary change between the two initial genotypes: *C*_*ab*_ = max(|Δ*E*_*a*_|; |Δ*E*_*b*_|)−|Δ*E*_*a*_ −Δ*E*_*b*_|, where Δ*E*_*a*_ and Δ*E*_*b*_ are the variations in expression between the evolved lineages *a* and *b* and their respective ancestors. *C*_*ab*_ is maximum when both lineages evolved in the same way, zero when one of the lineages did not evolve, and negative when both lineages evolved in opposite directions (Sup. Fig. 2). Convergent evolution scores were calculated between all combinations of evolutionary lineages and averaged out. Convergent evolution of gene expression plasticity (difference in gene expression between 23°C and 17°C) was also studied with the same index, Δ*E* then representing the difference in plasticity between evolved lineages and their ancestors.

#### Differential expression and enrichment

The statistical support for differential gene expression was estimated with the `DESeq2 package` for R (Love et al. 2014) from the uncorrected raw counts. The fixed-effect linear model was *G* + *S* + *E* + *L* + (*G* × *E*) + (*G* × *S*) + (*S* × *E*). The intercept for the “selection” factor was set to the ancestor, so that the effects of the selection regime directly quantify the gene expression evolution. We used a false discovery rate threshold of 0.05 associated with a minimal fold-change of 2 (i.e., | log_2_(*FC*) | ≥ 1) to build lists of differentially-expressed genes for each of the fixed effects. When more than two factor levels were available (Selection effect), DEG lists were merged (i.e., Selection DEGs were thus differentially-expressed in one or several selection regimes). Gene ontology enrichment was performed with topGO (Alexa and Rahnenfuhrer 2021); we used Fisher tests for the enrichment combined with the “weight” algorithm from topGO (Alexa et al. 2006). For completeness, we report lists of differentially-expressed genes obtained with a mixed-effect model (with the limma/dream procedure, Ritchie et al. (2015) and Hoffman and Roussos (2020)) in which Batch and Lineage were considered as random factors.

## Results

### Sources of transcriptomic variation

In this work, we propose to treat gene expression (measured as the logarithm of the RNAseq counts normalized and filtered as described in the Methods section) as quantitative multivariate phenotypic measurements. The logarithmic transformation improved to a large extent the skew and overdispersion in the distribution of gene expression, although a slightly negative mean-variance correlation remained (Sup. Fig. 1A). Average gene expressions at 17°C and 23°C were correlated to a large extent (*r* = 0:98, Sup. Fig. 1B), the difference between 23°C and 17°C was considered as a measurement of gene expression plasticity. Gene expression was well-correlated between both genotypes (*r* = 0:94, Sup. Fig. 1C), and interestingly, the residual variance (i.e., gene expression noise between biological replicates) was also similar between genotypes (*r* = 0:94, Sup. Fig. 1D). Plasticity was largely independent from gene expression (Sup. Fig. 1E), and was correlated between genotypes (*r* = 64, Sup. Fig. 1F)).

We first quantified the relative effect of experimental factors (genotype G: MGGP01 or MGGP44; selection regime S: Stable 17°C, Stable 23°C, or Fluctuating; and test temperature E, 17°C or 23°C) on gene expression. As the test temperature was the manipulated environment, from a plasticity perspective, genotype and selection measure changes in the elevation of the reaction norm, while temperature measures the slope. The variation in the dataset was structured by the differences between genotypes first, then by the plastic response to temperature, and finally by the evolutionary (“Selection”) effect (Fig. 2).

**Figure 2.**
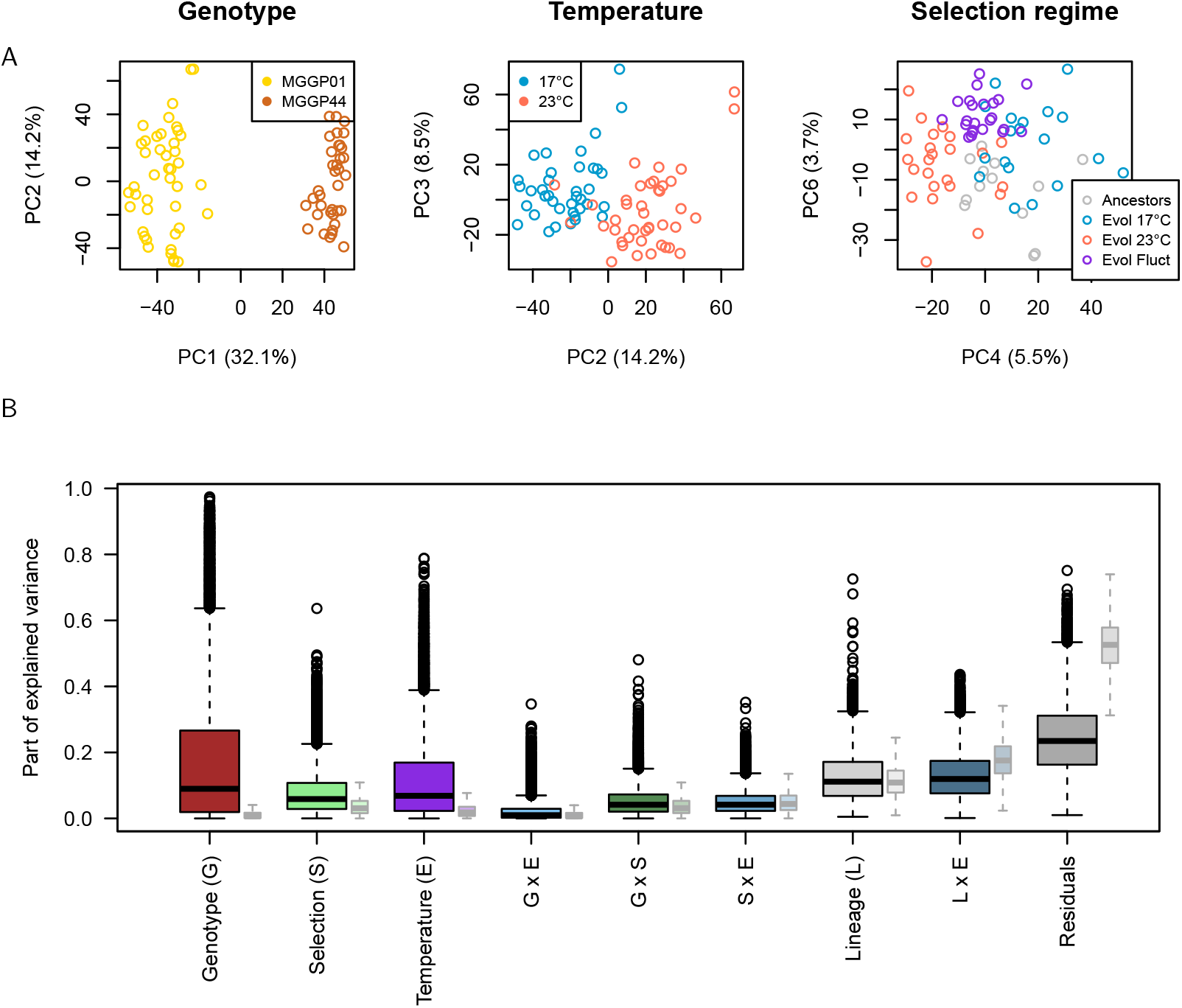
Sources of gene expression variation. A: Principal component analysis. PC1 catches the effect of the genotype, while PC2 (and, to a lesser extent, PC3) capture the effect of temperature (i.e., plasticity). Evolution is discriminated in PC4 and PC6. B: Distribution of the relative variance explained by experimental factors over all genes. Smaller boxes (control) display the same analysis in a column-permuted dataset.

When decomposing the variance in gene expression, we considered two layers of stochasticity: the “Lineage” effect captured the differences between evolutionary lineages submitted to the same selection regime (thus, genetic drift and/or mutational stochasticity), while the measurement noise (differences among biological replicates) fell into the residuals. For most genes, measurement noise was quite limited (median *<* 30% of the total variance), showing a remarkable reproducibility of the RNAseq procedure. The variance that was attributed to the selection effect (i.e., reproducible evolution) was slightly lower than the variance attributed to the lineage effect (non-reproducible evolution); note however that the lineage effect in the data was not larger than in a permuted dataset, as the experiment was not designed to study lineage-specific evolution at the scale of the transcriptome. We will thus focus on reproducible evolution, i.e. convergent evolution of gene expression between both initial genotypes submitted to the same selection regime, averaged over the triplicated evolutionary lineages.

We performed a differential analysis to assess the functional overlap between the genes involved in Genotype, Selection, and Environmental factors (Sup. Table 1). We listed differentially-expressed genes (DEGs) for Genotype, Lineage and Selection regime, Temperature, and their interactions, and we obtained 1462 DEGs (for a 5% false discrovery rate) for the Genotype effect, 638 DEGs for the Temperature effect (plastic genes), and 1732 DEGs for the Selection effect. For 319 genes, plasticity differed significantly between the genotypes (G×E interactions). Lists of differentially-expressed genes are provided as supplementary data. The functional annotation was substantially different for genes differing in average expression (Genotype effect) and genes differing in plasticity (G × E effect) (Sup. Fig. 3). While the average expression changes between genotypes was mostly associated with central metabolism and membrane transport, plasticity was associated with genes involved in specific biosynthetic pathways (glutathione, thiamine, and steroid) pathways. Interestingly, there was also no overlap between the function of universally plastic genes and genes differing in plasticity among genotypes (G × E). Genes that changed expression in the different selection treatments were associated with ceramide and glycogen pathways, as well as with epigenetic-related mechanisms (DNA and histones). This result suggests a relative independence between the genetic architecture of the differences between genotypes, sensitivity to temperature, and genes that are susceptible to adapt to new environmental challenges.

### Transcriptome adaptation involves massive convergent changes

Gene expression can be affected by both genetic and environmental (temperature) effects; it is traditional in quantitative genetics to measure the environmental effects as a linear reaction norm (variation of the trait across environments), which is defined by its slope (measuring phenotypic plasticity) and its intercept (the environment-independent expression). Here, we will follow the same logic; evolution of the intercept (the mid-point between the expression at two test temperatures) will be the topic of this section, while the evolution of the plastic slope (sensitivity to temperature) will be analysed in the next section.

Evolutionary change in the mean expression was substantial for some genes: in average, in a lineage, about 4.9% of the genes had their expression changed (in any direction) by more than one log2 unit, and 0.7% of the genes changed by 2 or more (Sup. Fig. 4). The fraction of expression-evolved genes was essentially identical between genotypes (*>* 1 log2-fold change: 5.2% in the background MGGP01, 4.6% in the background MGGP44). Evolutionary regimes were also very similar in terms of number of genes and magnitude of gene expression evolution. Evolution was not driving the populations in random directions, as gene expression change was substantially correlated in the two independent genotypes (fig. 3A). A multivariate redundancy analysis (RDA) constrained on the lineage (equivalent to a PCA from which environmental effects were removed) clearly identified an ancestor vs. evolved axis (RDA2) and a selection-specific axis (RDA3), fig. 3B. We considered this convergent evolution between genotypes as a signature of adaptive evolution.

**Figure 3.**
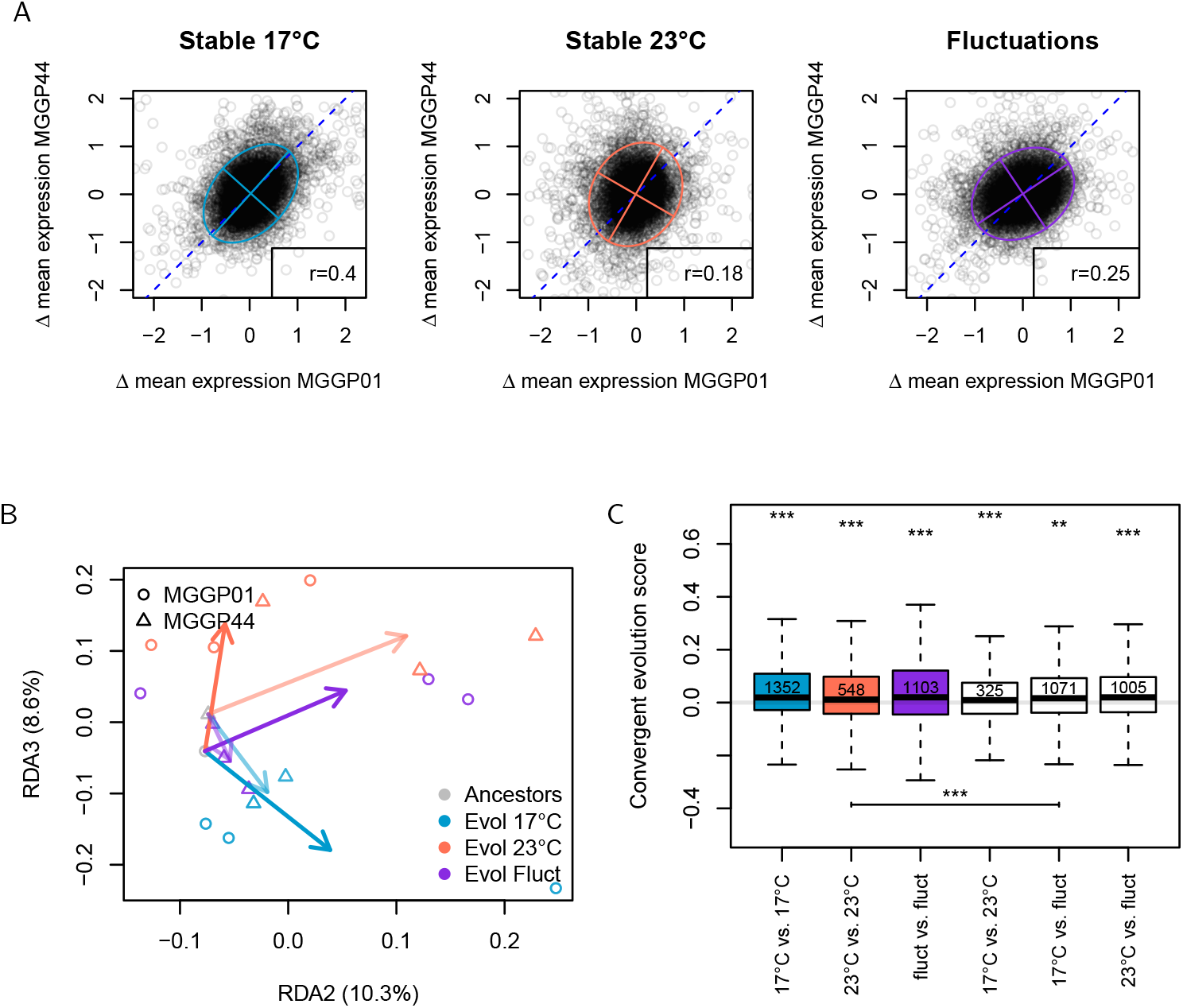
Convergent evolution of average gene expression in the three selection regimes. A: distribution of the change in gene expression in both genetic backgrounds. “Δ mean expression” stands for the difference in gene expression between the evolved populations and their ancestor, averaged over lineages (evolutionary replicates), biological replicates, and temperatures. The diagonal reference line materializes convergent evolution (same evolutionary change in both genotypes). Ellipses illustrate the covariance (95% confidence ellipses, assuming bivariate normal distributions). B: Redundancy Analysis considering lineages and genotypes as a grouping factor (19 levels in total, including evolutionary replicates for each selection regime in both ancestral genotypes). The first axis (RDA1, not shown) separates MGGP01 from MGGP44, the second axis RDA2 separates ancestors and evolved lineages, the third axis RDA3 catches the effect of selection regime. C: distribution of the convergent evolution scores between genotypes. Boxplots represent the quartiles of the distribution, whiskers expand to 1.5 times the interquartile distance. Outliers are not shown (they would span well beyond the limits of the figure). Stars denote the results of statistical tests (mixed-effect linear model, *C*_*ab*_ = selection+gene with gene considered as a random effect, p-values were obtained with the R package multcomp (Hothorn et al. 2008) which includes multiple testing correction). Above the boxes are displayed the p-values corresponding to the null hypothesis 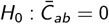, below the boxes are contrasted the convergent evolution of lineages from the same selection regime (colored boxes) vs. different regimes (white boxes) (*p* : 0:05 *<* ∗ *<* 0:01 *<* ∗∗ *<* 0:001 *<* ∗∗∗). The numbers within boxes correspond to the minimum number of genes that have to be removed before the loss of the statistical support for convergent evolution (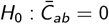, t-test, 5% p-value threshold).

As suggested by the large number of DEGs associated to the selection effect (1732 genes total, Sup. Fig. 3), the evolution at the transcriptome level was massive. This was reflected by the distribution of our convergent evolution scores, which were significantly biased towards positive values (fig. 3C). Convergent evolution occurred both between lineages that evolved in the same selection regime (e.g. Stable 17°C in MGGP01 vs. Stable 17°C in MGGP44) but also between lineages that evolved in different selection regimes (e.g. Stable 17°C in MGGP01 vs. Stable 23°C in MGGP44). The former can be interpreted as the effect of adaptation to the specific evolutionary conditions, while the later translates adaptation to the lab environment, irrespective of the selection regime. There were more convergent-evolved genes among lineages having evolved in the same selection regime (in average, 1001 vs. 800), and the difference was statistically supported (fig. 3C). We estimated the minimum number of convergent evolving genes by computing the statistical support for convergent evolution (p-value for *H*_0_ : 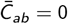) for reduced datasets, from which the genes with the maximal convergent evolution score were removed. There were at least 1352 genes which expression evolved in the same way in both genotypes at 17°C (i.e., one has to remove at least 1353 convergent genes from the data set to stop rejecting *H*_0_), 548 genes at 23°C, and 1103 genes in the fluctuating conditions, corresponding to 13%, 5%, and 11% of the transcriptome, respectively.

### Gene expression plasticity is not an adaptation to fluctuating conditions

Plasticity was quantified as the (log2) expression difference between 23°C and 17°C (positive plasticity stands for higher expression at 23°C), averaged across biological replicates. About 11.6% of the plasticity measures were beyond 1 log-fold-change, and 2.1% were above 2 log-fold-changes; the median log-fold-change was about 0.35.

Surprisingly, gene expression plasticity changed mostly in lineages that evolved at 17°C, and less in other lineages (fig. 4). The reasons why plasticity evolved more at 17°C are unclear, and explanations could fall along two groups of hypotheses: (i) non-adaptive hypotheses: evolution at 17°C is less predictable and lineages diverged more than in the other experimental conditions (because of genetic drift, genetic draft, mutation bias, or specific but unknown selection features), or (ii) adaptive hypotheses: the new selection optimum was farther away or directional selection was stronger. The correlation between convergent evolution of gene expression plasticity in both genetic backgrounds appears to be positive in all three selection regimes (fig. 4C), and approximately twice as large at 17°C compared to 23°C and fluctuating regimes (*r* = 0:26 vs. *r* = 0:14 and *r* = 0:13, respectively). Convergent evolution of plasticity was substantial (fig. 4B), but the statistical support for selection-specific evolution was reduced; the multivariate analysis could identify a clear ancestor-evolved axis (PC3 in 4D), but PC4 could only isolate the stable 17°C regime, while stable 23°C and fluctuating were mixed up. Overall, these observations clearly support the adaptive hypothesis (ii): plasticity evolved more at 17°C, and this evolution is consistent across independent evolutionary replicates. In contrast, plasticity also evolved in Stable 23°C and Fluctuating regimes, but to a lesser extent, and there is little evidence that this evolution was regime-specific.

**Figure 4.**
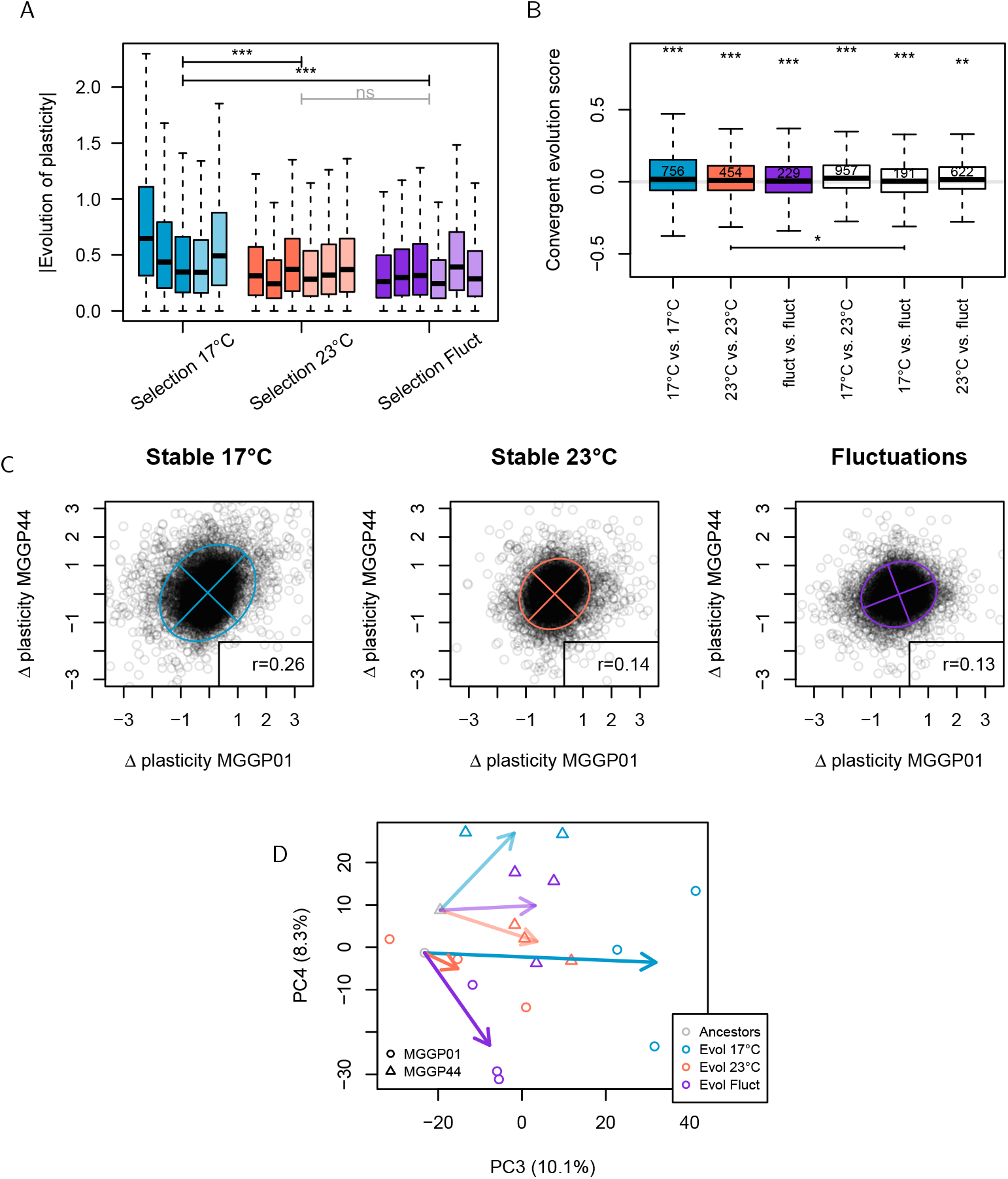
A: Distribution of the evolution of plasticity |Δplasticity| in the three experimental selection regimes. Dark colors stand for the genotype MGGP01, light colors for the genotype MGGP44. Boxplots indicate the median, 1st, and 3rd quartiles, whiskers expand to 1.5 times the interquartile distance. Outliers are not shown (they would span beyond the plotting area). The statistical differences among selection regimes were estimated by a generalized linear model (log gaussian link). B: Distribution of convergent evolution scores when comparing the same selection regime (colored boxes) and different selection regimes (white boxes), same legend as in Figure 3. C: Distribution of the evolution of plasticity in both genotypes for each selection regime. D: Principal Component Analysis considering plasticity (averaged over biological replicates) in evolved and ancestral lineages. Arrows represent the average evolution (from each ancestor to average evolved lineages). PC1 and PC2 (not shown) were not associated with lineage evolution. PC3 tends to separate ancestors to evolved lineages (adaptation to the lab environment), PC4 is associated with selection-specific evolution of plasticity, which repeatability across ancestral genotypes is much less convincing than for average gene expression.

There was very little to no association between the plasticity in the ancestral genotypes and the evolution of plasticity itself (Sup. Fig. 5). In other words, a gene that was ancestrally plastic was not more keen to evolve towards more or less plasticity than a gene that was initially not plastic. Correlations were of the same low magnitude (not shown) when considering the absolute value of plasticity (i.e., considering plastic vs non-plastic genes irrespective to the direction of plasticity). This confirms that plastic genes in the ancestor are not more not less evolvable than non-plastic genes, both for the intercept and the slope of the reaction norm.

### Environmental fluctuations drive gene expression evolution

Plasticity could evolve by changing the gene expression at 17°C, at 23°C, or both. We quantified the relative contributions of temperature-dependent gene expressions in the evolution of plasticity for the three selection regimes. Assuming that the final gene expression at a specific stable temperature could be considered as a proxy for the optimal gene expression at that temperature, we quantified for each gene how far evolution has driven gene expression towards this optimum (fig. 5A). In stable regimes, the evolutionary pattern was consistent with the intuitive idea that fluctuating selection was somehow intermediate in terms of evolutionary pressures, as lineages evolved under fluctuations were always intermediate between the optimum and the ancestor. Interestingly, the lineages that were never submitted to one of the temperatures also progressed towards the optimum at this temperature; this reflects the convergent gene expression changes across selection regimes (adaptation to the lab environment). Fluctuating populations may have not reached the optimum for several reasons, either because the need to adapt to several environments at the same time constrains evolution, or because they have simply been exposed less frequently (half the total time) to the selective environment. The same analysis considering evolution of the slope of the reaction norm (“plasticity” in fig. 5A) also features a progress for the fluctuating populations towards the theoretical best reaction norm compared to the stable selection populations, which remained close to the ancestral plasticity. Yet, the progress was of low amplitude, as if evolution of plasticity was substantially slower than evolution of gene expression. These results were robust to the details of the statistical and bioinformatics procedure, including count normalization and batch effect correction methods (Sup. Fig. 6). The reported patterns were still observable in absence of normalization or batch effect correction, demonstrating that they were not a by-product of the data treatment.

**Figure 5.**
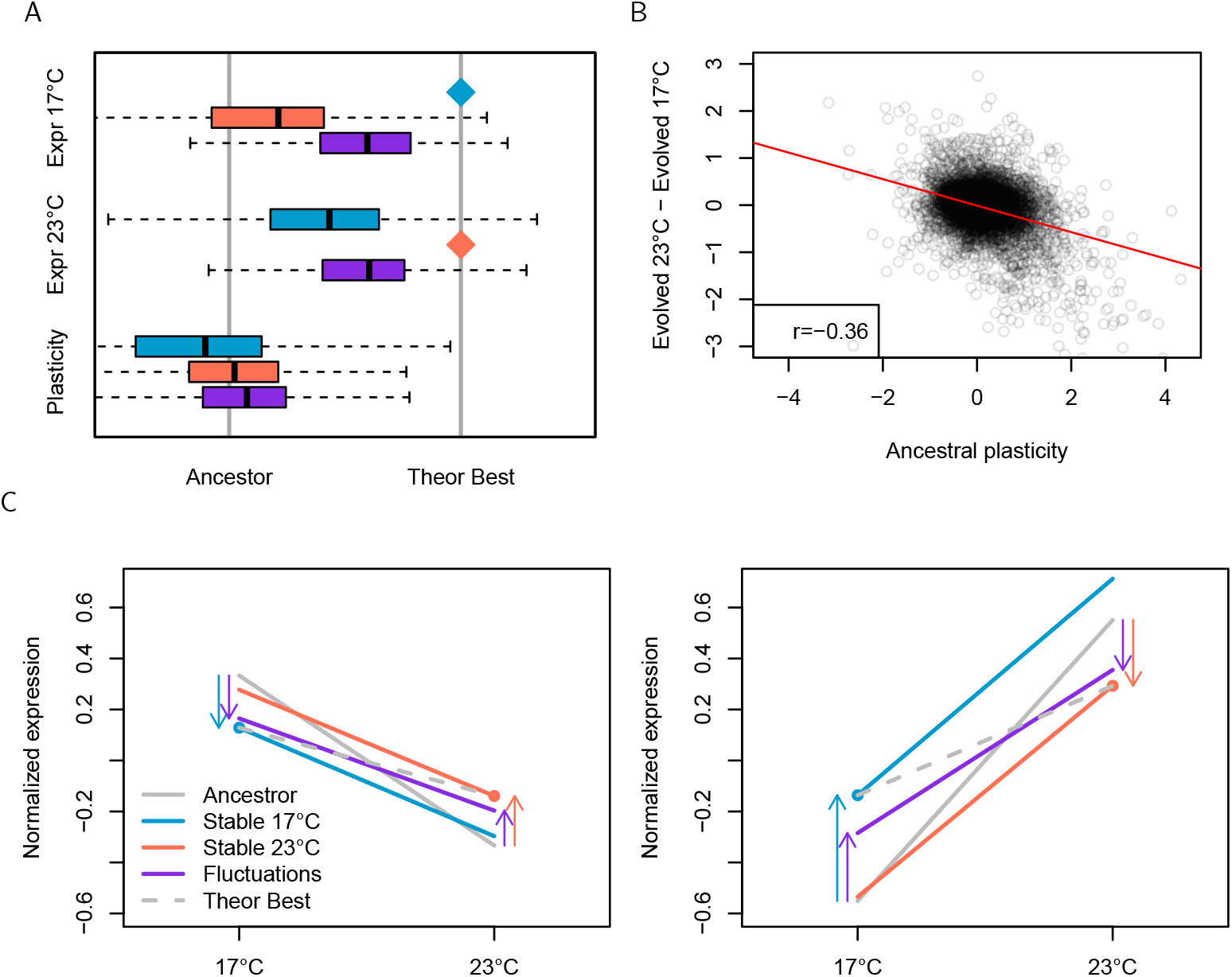
Mode of evolution for gene expression plasticity. A: Distribution of the evolution of gene expression relative to the distance to the putative optimum (“theoretical best”). For the expression at 17°C, the theoretical best is the mean expression in lineages having evolved at constant 17°C (blue diamond); for expression at 23°C, the optimum is the mean expression of lineages having evolved at 23°C (red diamond). For plasticity (=Expression at 23°C Expression at 17°C), the theoretical best is the difference between the two previous optima. In order to limit the measurement noise, only genes that had to evolve by *>* 1 fold-change are displayed in the figure. B: Negative correlation between the ancestral plasticity (averaged over both genotypes) and the difference between evolved stable regimes (averaged over both genotypes and both temperatures). C: Average reaction norms before (“Ancestor”) and after evolution, for genes displaying negative plasticity (left) and positive plasticity (right) in the ancestors (quantiles 0-10%, and 90-100%, respectively). The “theoretical best” plasticity is shown as a hyphenated line. Arrows indicate opportunity for direct selection.

Strikingly, transcriptomes evolved in a general direction opposite to the ancestral plasticity (fig. 5B), which is not expected if the ancestral plasticity was adaptive. We illustrated the evolution of the reaction norm (slope and intercept) in both genotypes by calculating the average reaction norm (normalized around the average gene expression of the genes) for subsets of genes displaying negative and positive plasticity in the ancestral genotypes (fig. 5C). This analysis confirms that extreme initial plasticity tends to fade out. Since there was no general change in transcrimptome plasticity (except in the Stable 17°C selection regime), the loss of initially-extreme plasticity had to be compensated by an increase in plasticity for some genes that were not very plastic in the ancestors. We would like to highlight three interesting observations that are meaningful in terms of transcriptome evolution: (i) in both environments, the fluctuating populations evolve in the same direction (but to a smaller distance) than the stable populations for each environment. This seems to confirm that, at this whole transcriptome scale, adaptation to fluctuations seems equivalent to adaptation to both environments; (ii) plasticity fades out faster in the fluctuating regime than in stable regimes, reinforcing the idea that for initially plastic genes, the loss of plasticity in fluctuating environments is associated with adaptation. However, this decrease in plasticity is not fast enough to catch up with the “theoretical best” plasticity, which reaction norm is always flatter; (iii) for this subset of initially plastic genes, populations in stable selection regimes experience little gene expression evolution (in average) in the environment they have never experienced. This suggests that genetic correlation between gene expression in both temperature environments may not be a major constraint in terms of plasticity evolution.

## Discussion

Here, we applied a quantitative approach to the analysis of the transcriptome during experimental evolution in different selection regimes. We showed that gene expression evolved substantially during the experiment, and the analysis of convergent evolution from independent genotypes suggests that evolution is at least partly driven by adaptation and affects a substantial fraction of the transcriptome. Furthermore, gene expression plasticity (temperature-dependent expression) also evolved substantially, but contrary to our expectation, plasticity did not evolve more in fluctuating conditions than in stable conditions.

### Transcriptome adaptation is massively pleiotropic

Gene expression change is a major contributor to adaptation (Fraser 2013). A substantial part of the transcriptome is susceptible to evolve among close species (e.g., more than 2000 genes in Drosophila, Rifkin et al. (2003), and more than 3000 in Saccharomyces Li and Fay (2017))). Temperature is a prominent environmental factor, and is often suspected to be a major driver of climatic adaptation (Clarke 2003); about 3% of the transcriptome covaries with the latitude of the sampled populations in *Drosophila melanogaster* (Juneja et al. 2016), 5% in *Drosophila subobscura* (Porcelli et al. 2016), and up to 9% in the fish *Fundulus heteroclitus* (Dayan et al. 2015). Adaptation to temperature can be fast; multiple experimental evolution assays have demonstrated a widespread transcriptomic evolutionary response to temperature change, e.g. 5% of the transcriptome in *Escherichia coli* (Riehle et al. 2005), 7% in *Drosophila pseudoobscura* (Laayouni et al. 2007), and up to 32% in the earlier analysis of this experiment in *Z. tritici* (Jallet et al. 2020). Finding several hundred genes which expression evolved consistently as a response to temperature is thus consistent with the literature.

Phenotypic measurements in evolve and resequence experiments are typically affected by spurious statistical correlations due to shared comparison references, as evolutionary change is defined by the difference between evolved lineages and their shared ancestor. For instance, it is tempting to interpret convergent gene expression change in evolutionary replicates as evidence for adaptation, while at least part of the correlation emerges as a consequence of subtracting the same data point (the ancestral expression) to both observations. Correcting this bias is technically difficult; it is sometimes possible to support the significance of biological effects by contrasting the observed (biased + biological effects) correlation to resampled data (bias only), as in e.g. Ghalambor et al. (2015). Here, we exploited the fact that we had two independent ancestors, and whenever necessary, we estimated convergent evolution from each ancestor (and not between evolutionary lineages diverged from the same ancestor). This approach remains substantially underpowered, because it ignores any genotype-specific or lineage-specific evolution, but it ensures that the observed evolutionary trends are not statistically biased.

Convergent evolution could be attributed to two components: adaptation to the lab environment (the same genes evolved in all selection regimes), and specific adaptation to each environment (stable or fluctuating temperatures). Both contributions were statistically supported, but we estimated that about 800 genes evolved in a convergent way between regimes, and about 1000 within each selection regime, suggesting that only ∼200 genes were evolving as a response to each temperature regime. We found little evidence for specific signatures for stable vs. fluctuating regimes.

The fact that expression changed for hundreds of genes as a consequence of adaptation cannot be interpreted as evidence for positive selection on all of the genes’ expression. Indeed, the number of causal mutations that could be fixed during this experiment is orders of magnitude lower than the number of genes which expression (or plasticity) changed. The number of beneficial mutations (full or partial selective sweeps) was probably around *n* ∼ 3 (for a 20% fitness gain) to *n* ∼ 6 (for a 50% fitness gain) (Appendix 1): there was not enough time for many mutations of small effect to segregate, and more mutations of large effect would have induced larger fitness gains. This necessarily implies that the mutant haplotypes that fixed during the experiment affected the expression of hundreds of genes each. Two theoretical hypotheses could explain such an observation: (i) mutations are highly pleiotropic and affect multiple genes through cascading regulatory effects, or (ii) beneficial variants were genetically linked with many neutral or slightly deleterious mutations that were driven to high frequencies by the selective sweeps. We believe that hypothesis (ii) is rather incompatible with our observations, given that we inferred more than 1000 genes convergently evolving between genotypes. Accumulating about 1000 independent mutations in 225 generations requires more than 4 cis-regulatory mutations per cell division, which seems high when compared to mutation rate estimates in this species (Habig et al. 2021); conclusive evidence would be brought by DNA resequencing in evolved lines. More convincingly, genetic draft or linkage disequilibrium could indeed change the expression of multiple neutral genes along with the fixation of an adaptive allele, but there is no reason why convergent changes would occur in independent evolutionary lineages. We thus deem it likely that in our experiment, adaptation was driven by the fixation of mutations that were both beneficial and highly pleiotropic, an observation that mirrors recent large-scale transcriptomic studies (Rennison and Peichel 2022; Ruelens et al. 2023; Koch et al. 2025).

Understanding how the expression of so many genes could evolve during short-term adaptation is challenging. Pleiotropic regulatory mutations have the potential to affect multiple genes, but pleiotropy is generally associated with deleterious effects (Vande Zande et al. 2022). One thus has to assume that single mutations in our experimental setup can have beneficial effects large enough to overwhelm hundreds of non-adaptive changes — in other words, most gene expression changes have to be neutral. Yet, the expression of many genes is generally reported to be under natural selection (e.g. 22% of temperature-related genes according to Whitehead and Crawford (2006)), most being under stabilizing selection (Gilad et al. 2006; McGuigan et al. 2014). A more controversial alternative is that gene networks may be organized in such a way that mutations trigger a coherent set of changes in the expression of multiple genes, maximizing the chances to generate non-antagonistic changes in the cell physiology. In other words, the genetic architecture of complex developmental systems could have evolved to favor pleiotropic effects that have synergistic consequences on fitness. Even if theoretically possible (Jones et al. 2014; Petit et al. 2023), and despite some empirical support (Houle et al. 2017; Collet et al. 2018), this hypothesis remains speculative, and restricted to the cases where future evolutionary challenges are similar to the past ones (possible for temperature fluctuations, much less for e.g. plant or animal domestication).

### Fluctuating selection is not associated with an increase in transcriptome plasticity

Coping with new environments does not solely rely on a genetic change of the gene expression, as gene expression plasticity is expected to play a key role in adaptation (López-Maury et al. 2008). Yet, experimental evolution focusing on gene expression plasticity remains scarce. In a one-year experiment with fluctuating salinity in the green algae *Dunaleilla salina*, Leung et al. (2020) showed that morphological (cell shape) plasticity decreased when the environment (salinity) was less predictable, as theoretically expected. This trend was paralleled with some plasticity loss in gene expression and epigenetic marks (Leung et al. 2023), confirming that this response to environmental fluctuations involved multiple organization levels. Plasticity thus evolved in the direction predicted by theoretical models in *Dunaleilla* (less plasticity in unpredictable environments). This sounds qualitatively different from our results in *Zymoseptoria*, but the selection regimes were also substantially different (predictable vs. unpredictable salinity fluctuations in *Dunaleilla*, compared to fluctuating vs. stable temperatures in our experiment). In a different context, Mallard et al. (2020) tracked the evolution of gene expression plasticity in *Drosophila* in cold (15°C) vs hot (23°C) environments. They noticed a general gain of plasticity at high temperatures (which we did not observe, as our observations rather support a gain of plasticity at low temperatures), and more evolution in plasticity than in mean gene expression; the lack of fluctuating regime compared to our design hinders further comparison. Closer to our experimental setup (but on different environmental factors), in Drosophila, Yampolsky et al. (2012) did not detect any evolution of gene expression plasticity in stable vs. fluctuating exposure to ethanol, while Huang and Agrawal (2016) observed counter-gradient evolution (i.e., maladaptive initial plasticity) in constant vs. fluctuating diets. Overall, so far, experimental evolution arguably failed at providing consistent observations about gene expression plasticity. The underlying reasons (different species, different environmental factors, different kinds of fluctuations) remain largely unknown.

Preferential evolution of gene expression plasticity in the Stable 17°C selection regime came as a surprise, although it confirms the previous observation by Jallet et al. (2020) on a partial dataset from the same experiment. The phenomenon was observed in both genotypes, and involves a lot of previously non-plastic genes, that became temperature dependent during the experiment. It is worth noting that this evolution of plasticity was not similar to what was observed in the fluctuating regime (it was rather in the opposite direction), which discards the possibility of minor undesired temperature fluctuations in the 17°C regime. Overall, the hypothesis that plasticity evolves as an adaptation to changing environments is not supported by our experiment. Here, it rather seems that initially-plastic genes evolved towards less plasticity in average, especially in fluctuating conditions; this plasticity loss was then compensated by a gain in plasticity for other genes. There was some substantial convergent evolution for plasticity, which eliminates the possibility that its evolution was driven by genetic drift. This result is consistent with counter-gradient evolution targeting the genes that were the most plastic in the initial populations, suggesting that adaptation tends to cancel initial plasticity. Maladaptive plasticity is a rather common observation (Ghalambor et al. 2015; Huang and Agrawal 2016; Ho and Zhang 2018; Koch and Guillaume 2020), and offers a satisfactory explanation for the lack of general plastic response in the fluctuating environment. Indeed, fluctuating environments expose the fitness consequences of maladaptive plasticity, and are thus associated with a decrease in the slope of the reaction norms. However, the rationale for the presence of maladaptive plasticity in the initial genotypes that were sampled from wild populations remains to be explained, as it sounds extremely unlikely that wild populations may not be adapted to temperature fluctuations. Alternative explanations must involve some reversal of the optimal reaction norm in the lab, or some trade-offs in the plastic response to different environmental variables, which are all difficult to validate empirically. In any case, a better understanding of the evolutionary challenges raised by the need to adapt to warmer and/or variable temperature regimes remains central to predicting the consequences of global change on biodiversity.

Why convergent plasticity evolves in stable environments is not totally clear neither, because it suggests that gene expression in a environment that is never encountered may follow some repeatable trend. Genetic correlations across environments could explain such a pattern, but the observed genetic decorrelation between the evolution of the intercept and the slope of reaction norms does not support this hypothesis. Alternatively, plasticity could evolve as a response to fluctuations to environmental variables independent from temperature, such as cell density or nutrient concentration, which vary periodically as a consequence of the experimental design. Due to the large number of genes involved in adaptation to the lab environment (not selection-specific), it is tempting to propose that the effect of temperature on most of these genes might be the consequences of some form of pleiotropy, as a side effect of the response to some unmeasured fluctuating factors. Little is known about the modularity and genetic independence of the plastic response to different environmental variables, and alternative experimental setups, involving more than one environmental variables, would be necessary to confirm such speculations.

## Conclusion

We were able to evidence here some repeatable evolutionary trends from a whole transcriptome approach. Based on convergent evolution in independent evolutionary replicates, we showed that adaptation was associated to gene expression changes throughout the whole system: our results suggest that gene expression evolves at a global, system-wide scale, and not restricted to specific genes or small-scale modules. Contrary to our expectations based on classical theory, plasticity did not evolve more in fluctuating environments, and ideal reaction norms were shallower than the observed plasticity. These observations do not fit very well with known predictions from gene-centered (population genetics) nor trait-centered (quantitative genetics) approaches, which highlights the dire need for alternative interpretative theoretical frameworks in evolutionary systems biology.

## Data & Scripts

All RNA-seq transcriptome data generated in this study have been deposited in the NCBI BioProject PRJNA1414243.

The bioinformics pipeline (from raw data to counts) is available at https://github.com/genissellab/rna-evolve-reseq.

Normalized count data and analysis scripts are provided at https://github.com/lerouzic/Quantizymo.

## Acknowledgements

We thank Johann Confais for his help during experimental evolution.

## Appendix 1 number of selective sweeps

The number of selective sweeps involved in a given fitness gain in a given amount of time is constrained by the speed of adaptation (smaller effect mutations segregate slower). In non-recombining populations, clonal interference also limits the number of adaptive steps.

### From standard population genetics

In infinite, asexual, and haploid organisms, it takes *t* = *L*_0→1_ /*s* generations to evolve from frequency *p*_0_ to *p*_1_ for a mutation of fitness effect +*s*, with *L*_0→1_ = log[(*p*_1_(1 − *p*_0_))*=*(*p*_0_(1 − *p*_1_))] (equation 5.3.16 in Crow and Kimura 1970). If the experiment lasts *T* generations and the total fitness gain is *S*, as a first approximation, the number of independent mutations of equal effect *s* = *S/n* on fitness that could generate successive selective sweeps is of the order of 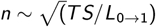. We estimated the total number of generations to be about *T* = 225, and preliminary estimates of the total fitness (growth rate) gains ranged between *S* = 0:2 and *S* = 0:5 depending on the lineages. If a selective sweep consists in raising from frequency *p*_0_ = 0:1 to *p*_1_ = 0:9, *L*_0→1_ ≃ 4:39, and *n* ranges from 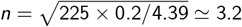 to 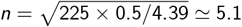 for fitness gains of *S* = 0:2 and *S* = 0:5, respectively.

### From fitness dynamic models

If the mutation rate is not too low, several beneficial mutations could in fact segregate together during adaptation, generating complex clonal interference patterns. A fitness dynamic model considering clonal interference (and epistasis for fitness, which we will disregard here) was proposed by Wiser et al. (2013), with a mean fitness increasing over time according to 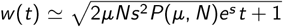, *s* being the effect of a single beneficial mutation, *—* representing the beneficial mutation rate, and 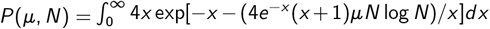 being proportional to the fixation time of a beneficial mutation. Average fitness *w* (*T* ) after *T* = 225 generations and the average number of sweeps (*w* (*T* ) − *w* (0))/*s* are represented in Appendix Fig 1A.

### From numerical simulations

Simple numerical simulations were run in populations of size *N* = 10^5^ for *T* = 225 non-overlapping generations, in which beneficial mutations improving fitness by a coefficient *s >* 0 occurred at a genomic rate *—*. Reproduction consisted in drawing *N* new clones from the parents with probabilities proportional to their fitness *w* = 1 + *ns*, where *n* was the number of beneficial mutations carried by each clone. Simulation results are displayed in Appendix Fig 1B. Simulations allowing for deleterious mutations were also run, with essentially identical results.

**Appendix Figure 1.**
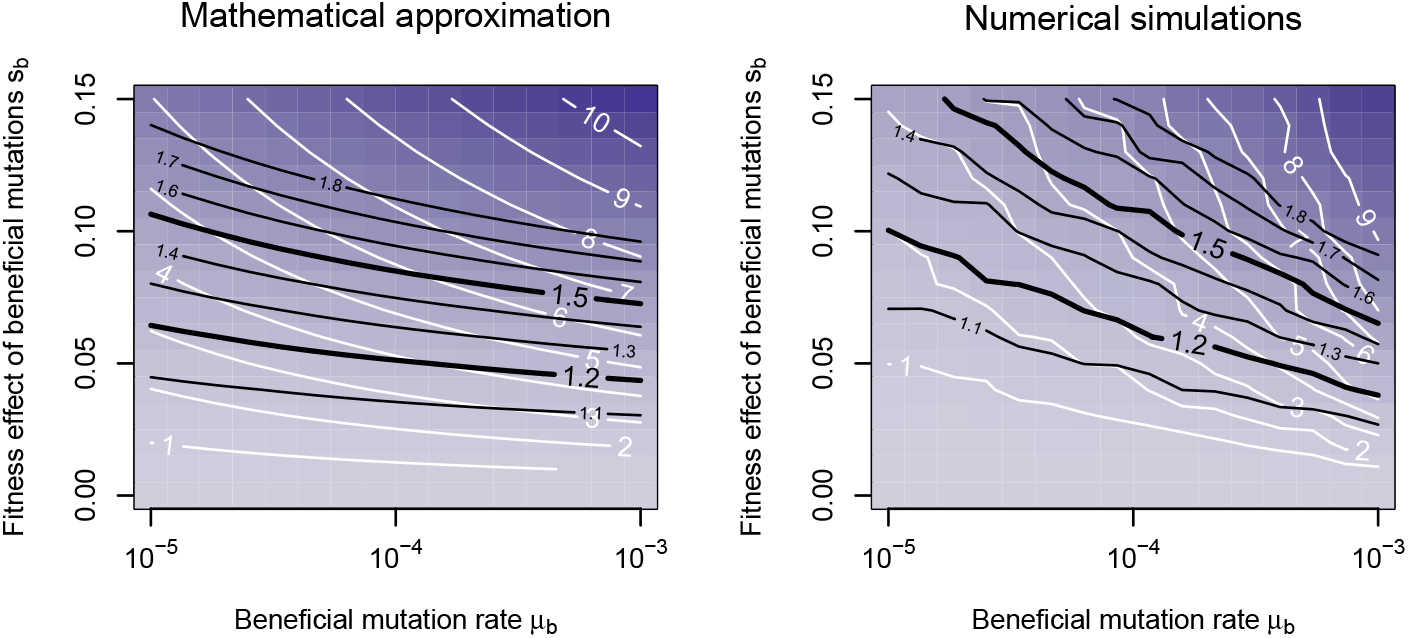
Average number of beneficial mutations (in white) and average fitness gain (contour lines in black and blue color shade) as a function of the rate of beneficial mutations and the effect of beneficial mutations. A: Mathematical approximation from Wiser et al. (2013), B: Numerical simulations. The range of fitness progress during experimental evolution (+20% to +50%) is highlighted; in this range, for realistic mutation rates, the average number of adaptive mutations remains between 2 and 6. The population size was set to *N* = 10^5^ in both models.

## References

Alexa, A. and J. Rahnenfuhrer (2021). topGO: Enrichment Analysis for Gene Ontology. R package version 2.46.0.

Alexa, A., J. Rahnenführer, and T. Lengauer (2006). “Improved scoring of functional groups from gene expression data by decorrelating GO graph structure”. In: Bioinformatics 22.13, pp. 1600–1607.

Bradshaw, A. D. (1965). “Evolutionary significance of phenotypic plasticity in plants”. In: Advances in genetics 13, pp. 115–155.

Chevin, L.-M., R. Lande, and G. M. Mace (2010). “Adaptation, plasticity, and extinction in a changing environment: towards a predictive theory”. In: PLoS biology 8.4, e1000357.

Clarke, A. (2003). “Costs and consequences of evolutionary temperature adaptation”. In: Trends in Ecology & Evolution 18.11, pp. 573–581.

Collet, J. M., K. McGuigan, S. L. Allen, S. F. Chenoweth, and M. W. Blows (2018). “Mutational pleiotropy and the strength of stabilizing selection within and between functional modules of gene expression”. In: Genetics 208.4, pp. 1601–1616.

Crow, J. F. and M. Kimura (1970). An introduction to population genetics theory. Harper & Row, Publishers, New York.

Dayan, D. I., D. L. Crawford, and M. F. Oleksiak (2015). “Phenotypic plasticity in gene expression contributes to divergence of locally adapted populations of F undulus heteroclitus”. In: Molecular ecology 24.13, pp. 3345–3359.

Dhar, R., R. Sägesser, C. Weikert, and A. Wagner (2013). “Yeast adapts to a changing stressful environment by evolving cross-protection and anticipatory gene regulation”. In: Molecular biology and evolution 30.3, pp. 573–588.

Fay, J. and P. Wittkopp (2008). “Evaluating the role of natural selection in the evolution of gene regulation”. In: Heredity 100.2, pp. 191–199.

Fraser, H. B. (2013). “Gene expression drives local adaptation in humans”. In: Genome Research 23.7, pp. 1089–1096. doi: 10.1101/gr.152710.112.

Garland, T. and M. R. Rose (2009). Experimental evolution: concepts, methods, and applications of selection experiments. Univ of California Press.

Ghalambor, C. K., K. L. Hoke, E. W. Ruell, E. K. Fischer, D. N. Reznick, and K. A. Hughes (2015). “Nonadaptive plasticity potentiates rapid adaptive evolution of gene expression in nature”. In: Nature 525.7569, pp. 372–375.

Gilad, Y., A. Oshlack, and S. A. Rifkin (2006). “Natural selection on gene expression”. In: TRENDS in Genetics 22.8, pp. 456–461.

Habig, M., C. Lorrain, A. Feurtey, J. Komluski, and E. H. Stukenbrock (2021). “Epigenetic modifications affect the rate of spontaneous mutations in a pathogenic fungus”. In: Nature Communications 12.1, p. 5869.

Ho, W.-C. and J. Zhang (2018). “Evolutionary adaptations to new environments generally reverse plastic phenotypic changes”. In: Nature communications 9.1, p. 350.

Hoffman, G. E. and P. Roussos (2020). “dream: Powerful differential expression analysis for repeated measures designs”. In: Bioinformatics. doi: 10.1093/bioinformatics/btaa687.

Hothorn, T., F. Bretz, and P. Westfall (2008). “Simultaneous Inference in General Parametric Models”. In: Biometrical Journal 50.3, pp. 346–363.

Houle, D., G. H. Bolstad, K. van der Linde, and T. F. Hansen (2017). “Mutation predicts 40 million years of fly wing evolution”. In: Nature 548.7668, pp. 447–450.

Huang, Y. and A. F. Agrawal (2016). “Experimental evolution of gene expression and plasticity in alternative selective regimes”. In: PLoS genetics 12.9, e1006336.

Hughes, B. S., A. J. Cullum, and A. F. Bennett (2007). “An experimental evolutionary study on adaptation to temporally fluctuating pH in Escherichia coli”. In: Physiological and Biochemical Zoology 80.4, pp. 406–421.

Jallet, A. J., A. Le Rouzic, and A. Genissel (2020). “Evolution and plasticity of the transcriptome under temperature fluctuations in the fungal plant pathogen Zymoseptoria tritici”. In: Frontiers in Microbiology 11, p. 573829.

Jones, A. G., R. Bürger, and S. J. Arnold (2014). “Epistasis and natural selection shape the mutational architecture of complex traits”. In: Nature communications 5.1, p. 3709.

Juneja, P., A. Quinn, and F. M. Jiggins (2016). “Latitudinal clines in gene expression and cis-regulatory element variation in Drosophila melanogaster”. In: Bmc Genomics 17.1, p. 981.

Kawecki, T. J., R. E. Lenski, D. Ebert, B. Hollis, I. Olivieri, and M. C. Whitlock (2012). “Experimental evolution”. In: Trends in ecology & evolution 27.10, pp. 547–560.

Ketola, T. and K. Saarinen (2015). “Experimental evolution in fluctuating environments: tolerance measurements at constant temperatures incorrectly predict the ability to tolerate fluctuating temperatures”. In: Journal of Evolutionary Biology 28.4, pp. 800–806.

Koch, E. L. and F. Guillaume (2020). “Restoring ancestral phenotypes is a general pattern in gene expression evolution during adaptation to new environments in Tribolium castaneum”. In: Molecular ecology 29.20, pp. 3938–3953.

Koch, E. L., C. Rocabert, C. Beeravolu Reddy, and F. Guillaume (2025). “Gene expression evolution is predicted by stronger indirect selection at more pleiotropic genes”. In: Evolution Letters, qraf039.

Laayouni, H., F. García-Franco, B. E. Chávez-Sandoval, V. Trotta, S. Beltran, M. Corominas, and M. Santos (2007). “Thermal evolution of gene expression profiles in Drosophila subobscura”. In: BMC evolutionary biology 7.1, p. 42.

Laland, K., T. Uller, M. Feldman, K. Sterelny, G. B. Müller, A. Moczek, E. Jablonka, J. Odling-Smee, G. A. Wray, H. E. Hoekstra, et al. (2014). “Does evolutionary theory need a rethink?” In: Nature 514.7521, pp. 161–164.

Lande, R. (2009). “Adaptation to an extraordinary environment by evolution of phenotypic plasticity and genetic assimilation”. In: Journal of evolutionary biology 22.7, pp. 1435–1446.

Lapalu, N. et al. (Apr. 2023). “Improved gene annotation of the fungal wheat pathogenZymoseptoria triticibased on combined Iso-Seq and RNA-Seq evidence”. In: bioRxiv. doi: 10.1101/2023.04.26.537486.

Lenski, R. E., M. R. Rose, S. C. Simpson, and S. C. Tadler (1991). “Long-term experimental evolution in Escherichia coli. I. Adaptation and divergence during 2,000 generations”. In: The American Naturalist 138.6, pp. 1315–1341.

Leroi, A. M., R. E. Lenski, and A. F. Bennett (1994). “Evolutionary adaptation to temperature. III. Adaptation of Escherichia coli to a temporally varying environment”. In: Evolution 48.4, pp. 1222–1229.

Leung, C., D. Grulois, L. Quadrana, and L.-M. Chevin (2023). “Phenotypic plasticity evolves at multiple biological levels in response to environmental predictability in a long-term experiment with a halotolerant microalga”. In: PLoS Biology 21.3, e3001895.

Leung, C., M. Rescan, D. Grulois, and L.-M. Chevin (2020). “Reduced phenotypic plasticity evolves in less predictable environments”. In: Ecology letters 23.11, pp. 1664–1672.

Lewens, T. (2019). “The Extended Evolutionary Synthesis: what is the debate about, and what might success for the extenders look like?” In: Biological Journal of the Linnean Society 127.4, pp. 707–721.

Li, X. C. and J. C. Fay (2017). “Cis-regulatory divergence in gene expression between two thermally divergent yeast species”. In: Genome biology and evolution 9.5, pp. 1120–1129.

Long, A., G. Liti, A. Luptak, and O. Tenaillon (2015). “Elucidating the molecular architecture of adaptation via evolve and resequence experiments”. In: Nature Reviews Genetics 16.10, pp. 567–582.

López-Maury, L., S. Marguerat, and J. Bähler (2008). “Tuning gene expression to changing environments: from rapid responses to evolutionary adaptation”. In: Nature Reviews Genetics 9.8, pp. 583–593. doi: 10.1038/nrg2398.

Love, M., S. Anders, and W. Huber (2014). “Differential analysis of count data–the DESeq2 package”. In: Genome Biol 15.550, pp. 10–1186.

Mallard, F., V. Nolte, and C. Schlötterer (2020). “The evolution of phenotypic plasticity in response to temperature stress”. In: Genome Biology and Evolution 12.12, pp. 2429–2440.

McGuigan, K., J. M. Collet, S. L. Allen, S. F. Chenoweth, and M. W. Blows (2014). “Pleiotropic mutations are subject to strong stabilizing selection”. In: Genetics 197.3, pp. 1051–1062.

Pennacchi, J. P., J. M. S. Lira, M. Rodrigues, F. H. S. Garcia, A. M. d. C. Mendonca, and J. P. R. A. D. Barbosa (2021). “A systemic approach to the quantification of the phenotypic plasticity of plant physiological traits: the multivariate plasticity index”. In: Journal of Experimental Botany 72.5, pp. 1864–1878.

Petit, A. J., J. Guez, and A. Le Rouzic (2023). “Correlated stabilizing selection shapes the topology of gene regulatory networks”. In: Genetics 224.2, iyad065.

Porcelli, D., A. M. Westram, M. Pascual, K. J. Gaston, R. K. Butlin, and R. R. Snook (2016). “Gene expression clines reveal local adaptation and associated trade-offs at a continental scale”. In: Scientific reports 6.1, p. 32975.

Price, P. D., D. H. Palmer Droguett, J. A. Taylor, D. W. Kim, E. S. Place, T. F. Rogers, J. E. Mank, C. R. Cooney, and A. E. Wright (2022). “Detecting signatures of selection on gene expression”. In: Nature Ecology & Evolution 6.7, pp. 1035–1045.

R Core Team (2021). R: A Language and Environment for Statistical Computing. R Foundation for Statistical Computing. Vienna, Austria.

Rennison, D. J. and C. L. Peichel (2022). “Pleiotropy facilitates parallel adaptation in sticklebacks”. In: Molecular ecology 31.5, pp. 1476–1486.

Riehle, M. M., A. F. Bennett, and A. D. Long (2005). “Changes in gene expression following high-temperature adaptation in experimentally evolved populations of E. coli”. In: Physiological and Biochemical Zoology 78.3, pp. 299–315.

Rifkin, S. A., J. Kim, and K. P. White (2003). “Evolution of gene expression in the Drosophila melanogaster subgroup”. In: Nature genetics 33.2, pp. 138–144.

Ritchie, M. E., B. Phipson, D. Wu, Y. Hu, C. W. Law, W. Shi, and G. K. Smyth (2015). “limma powers differential expression analyses for RNA-sequencing and microarray studies”. In: Nucleic Acids Research 43.7, e47. doi: 10.1093/nar/gkv007.

Ruelens, P., T. Wynands, and J. A. G. de Visser (2023). “Interaction between mutation type and gene pleiotropy drives parallel evolution in the laboratory”. In: Philosophical Transactions of the Royal Society B 378.1877, p. 20220051.

Schmalhausen, I. I. (1949). “Factors of evolution: the theory of stabilizing selection.” In.

Sommer, R. J. (2020). “Phenotypic plasticity: from theory and genetics to current and future challenges”. In: Genetics 215.1, pp. 1–13.

Spitze, K. and T. D. Sadler (1996). “Evolution of a generalist genotype: multivariate analysis of the adaptiveness of phenotypic plasticity”. In: The American Naturalist 148, S108–S123.

Van den Bergh, B., T. Swings, M. Fauvart, and J. Michiels (2018). “Experimental design, population dynamics, and diversity in microbial experimental evolution”. In: Microbiology and Molecular Biology Reviews 82.3, pp. 10–1128.

Vande Zande, P., M. S. Hill, and P. J. Wittkopp (2022). “Pleiotropic effects of trans-regulatory mutations on fitness and gene expression”. In: Science 377.6601, pp. 105–109.

Via, S. and R. Lande (1985). “Genotype-environment interaction and the evolution of phenotypic plasticity”. In: Evolution 39.3, pp. 505–522.

Whitehead, A. and D. L. Crawford (2006). “Neutral and adaptive variation in gene expression”. In: Proceedings of the National Academy of Sciences 103.14, pp. 5425–5430.

Wiser, M. J., N. Ribeck, and R. E. Lenski (2013). “Long-term dynamics of adaptation in asexual populations”. In: Science 342.6164, pp. 1364–1367.

Yampolsky, L. Y., G. V. Glazko, and J. D. Fry (2012). “Evolution of gene expression and expression plasticity in long-term experimental populations of Drosophila melanogaster maintained under constant and variable ethanol stress”. In: Molecular Ecology 21.17, pp. 4287–4299.

