## Supplementary figures 1 to 6, Sup Table 1 for "Adaptive evolution of transcriptome and transcriptomic plasticity in stable and fluctuating environments"

### Supplementary material for:
### Adaptive evolution of transcriptome and transcriptomic plasticity in stable and fluctuating environments

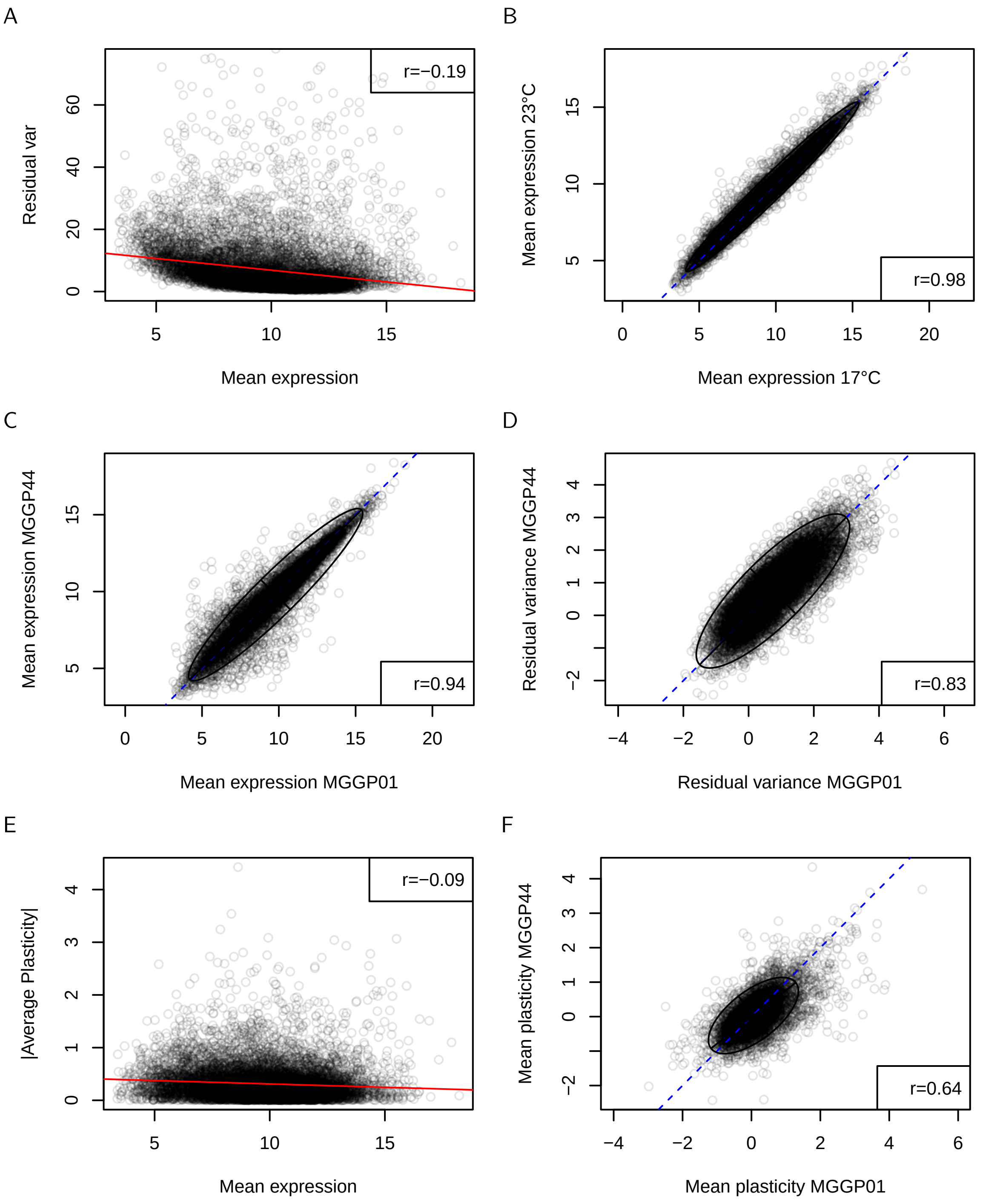


Supplementary Figure 1: Distribution of gene expressions in the dataset after bias correction. Expression is in log2(counts) units. A: grand mean expression vs. residual variance. B: expression at 17°C (all genotypes and selection regimes averaged) vs expression at 23°C. C to E: Comparison between genotypes of mean expression, residual variance, and mean plasticity. E: Mean expression vs. plasticity. C:

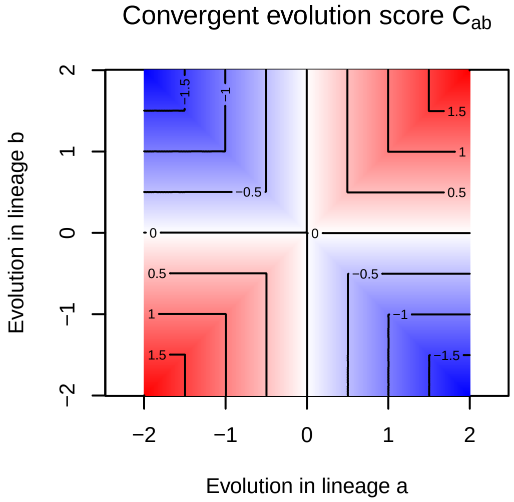


Supplementary Figure 2: The convergent evolution score $C_{ab}$ translates both the magnitude and the consistency of gene expression evolution between two evolutionary lineages $a$ and $b$. The figure illustrates that the maximum score (red) is obtained when the evolution is both large and consistent (either towards an increase or a decrease in the gene expression). When the expression evolves in a lineage but not in the other, the score is 0. Evolution in opposite directions lead to negative values (blue).

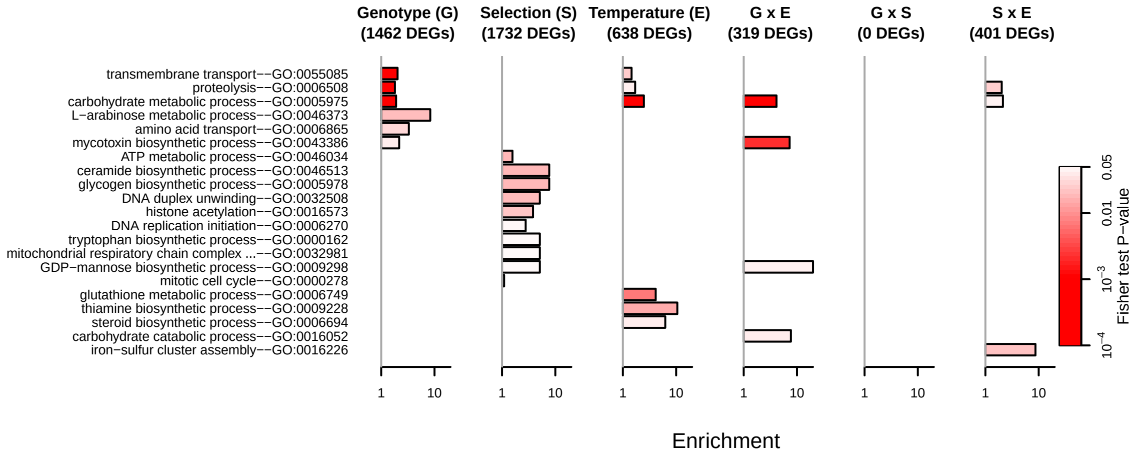


Supplementary Figure 3: Gene ontology enrichment for the sources of gene expression variation. Lists of DEGs were determined among genes which fold-changed was larger than 2 and a false discovery rate $< 0.2$. When more than two factor levels were involved (Selection regime level), lists of DEGs were merged. The figure represents only significant GO categories (p-value $< 0.05$) from Fisher exact tests, the color intensity increases when p-values decrease. Enrichment was calculated as the ratio between the observed number of significant DEGs and the expected number under the null hypothesis in each category.

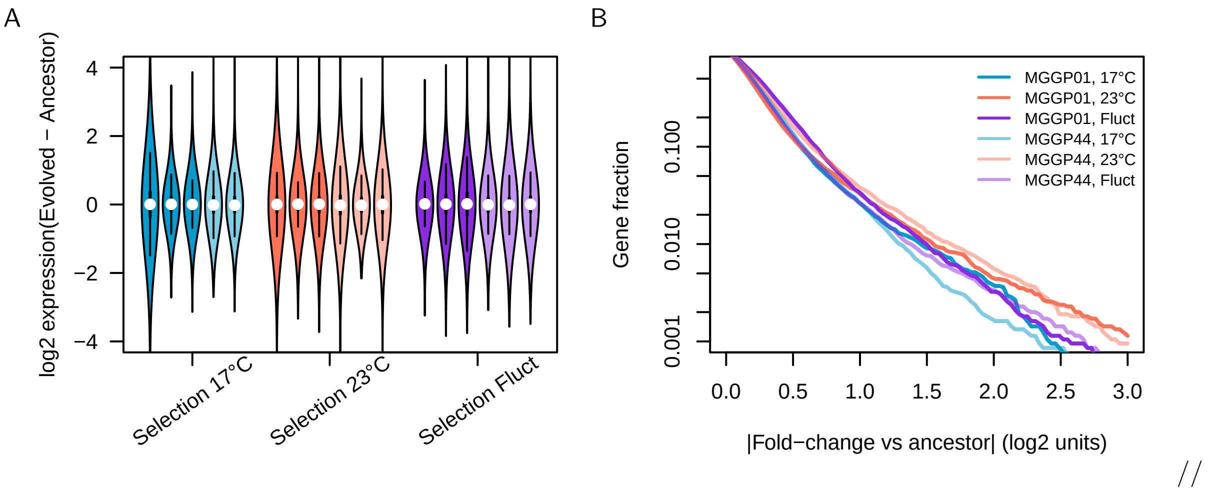


Supplementary Figure 4: Distribution of gene expression change during experimental evolution (averaged over the environments). A: Expression change (evolved - ancestor) in all evolutionary lineages (MGGP01 in dark colors, MGGP44 in light colors). B: Fraction of the genes with an absolute fold change > to the x axis, averaged by selection regime and genotype.

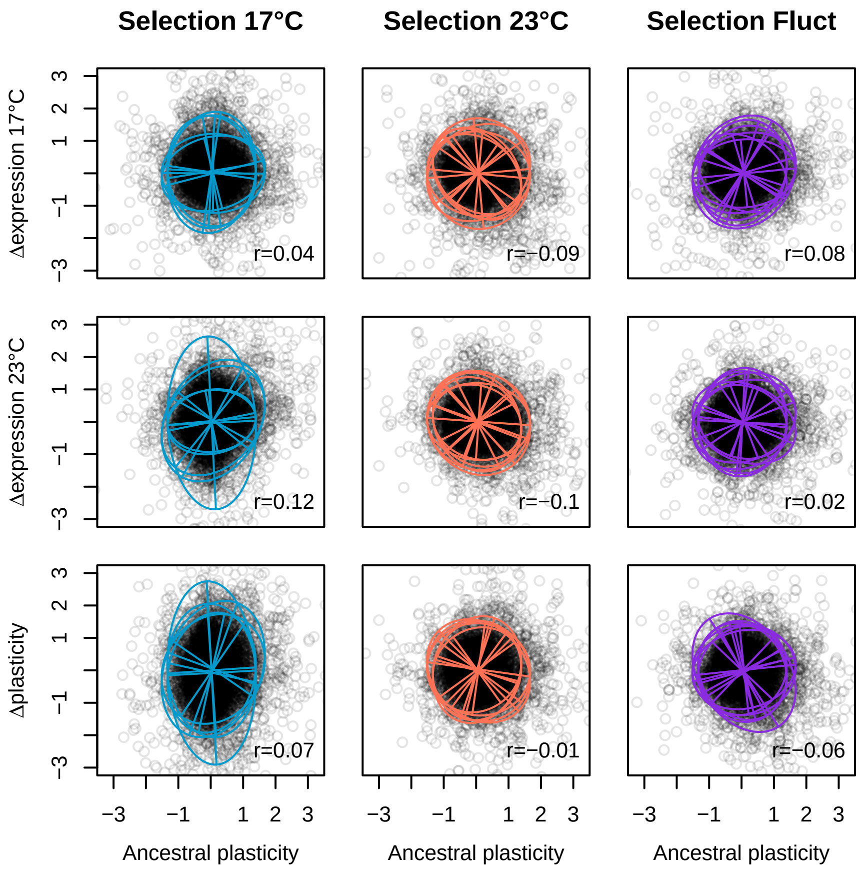


Supplementary Figure 5: Influence of the ancestral gene expression plasticity on the evolution of gene expression (first row: at 17°C, second row: at 23°C) and evolution of plasticity (bottom row). All replicates from the same selection regime were merged. In order to avoid spurious statistical associations, correlations were computed against the other ancestors (i.e., evolution of MGGP01 vs. plasticity in the ancestor of MGGP44 and vice versa).

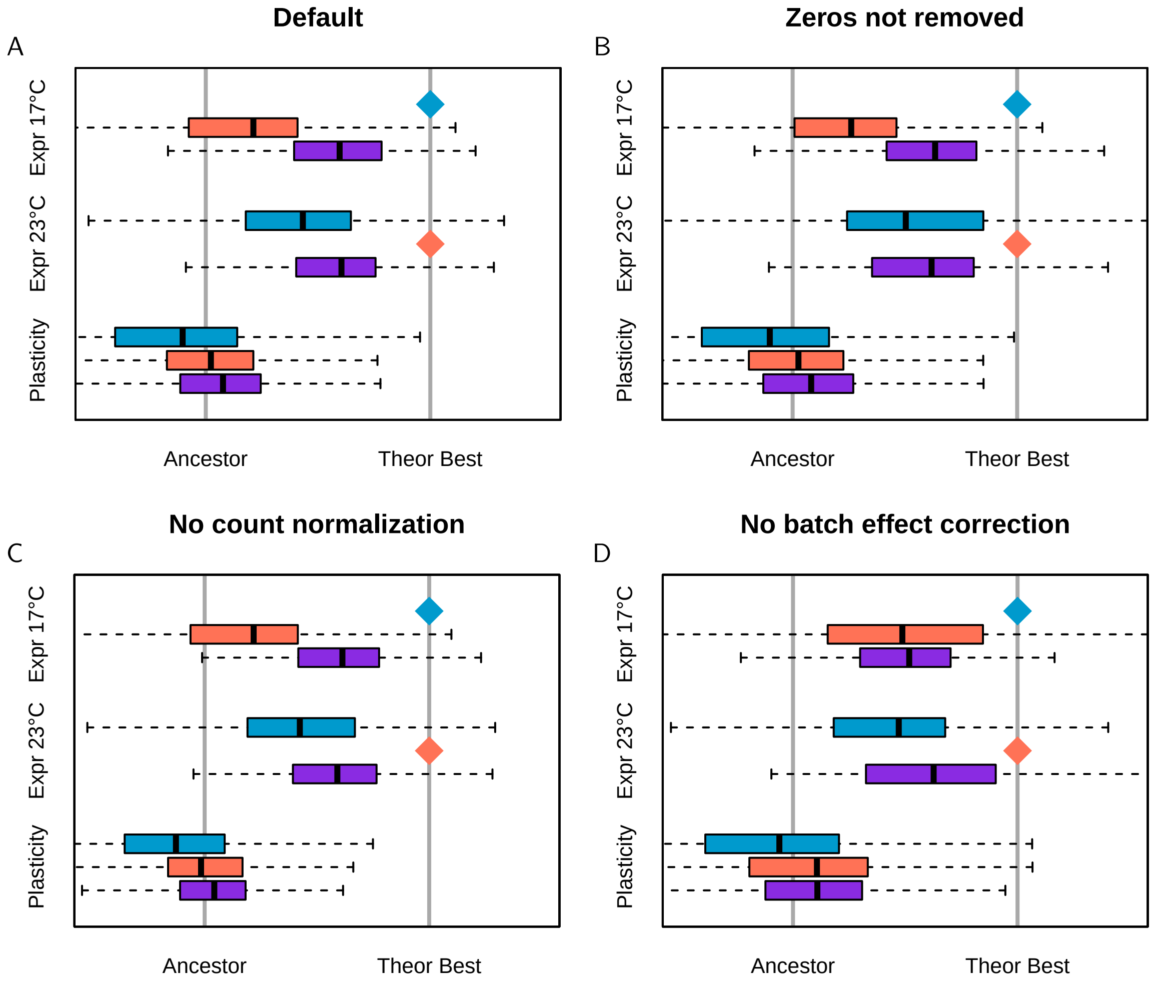


Supplementary Figure 6: Influence of data manipulation and curation on one of the main results (fig5A). A: Default; B: zero-expressed genes are kept; C: No count normalization; D: No batch effect correction.

| | Dream (random) | DESeq (fixed) |
|---|---|---|
| Genotype (G) | 808 | 1462 |
| Selection (S) | 47 | 1732 |
| Temperature (E) | 62 | 638 |
| Lineage (L) | 0 | 6670 |
| G $\times$ E | 436 | 319 |
| G $\times$ S | 17 | 0 |
| S $\times$ E | 269 | 401 |

Supplementary Table 1: Number of differentially-expressed genes (FDR $\leq$ 0.05 and log FC $\geq$ 1) for the different factors, with a mixed-effect model (dream) and a fixed-effect model (DESeq). In the mixed-effect model, Batch and Lineage were considered as random effects, which ensures that Temperature and Selection DEGs, respectively, cannot be attributed to sampling (at a substantial cost in terms of statistical power).
